# New gamma interferon (IFN-γ) algorithm for tuberculosis diagnosis in cynomolgus macaques

**DOI:** 10.1101/2024.04.04.588038

**Authors:** Saradee Warit, Suthirote Meesawat, Pattsarun Cheawchanlertfa, Nampueng Makhoa, Prapaporn Srilohasin, Machamon Kaewparuehaschai, Kirana Noradechanon, Areeya Pomcoke, Taratorn Kemthong, Therdsak Prammananan, Reka Kanitpun, Tanapat Palaga, Suchinda Malaivijitnond, Angkana Chaiprasert

## Abstract

Tuberculosis (TB) is the first infectious disease to be screened-out from specified pathogen-free cynomolgus macaques (*Macaca fascicularis*; Mf) using in human pharmaceutical testing. Being either latent or active stage after exposure to the *Mycobacterium tuberculosis* complex (MTBC), the monkey gamma-interferon release assay (mIGRA) was previously introduced for early TB detection in Mf. However, a high number of indeterminate cases were unexpectedly encountered. The main reasons were a mitogen positive control and an interpretation algorithm. A cohort of 316 Mf exposed to MTBC was tested of two positive mitogen controls [QFT-PHA and a mixture of ConcanavalinA and Pokeweed (ConA+PWM)], and 100 of 316 animals were selected and 26-month followed-up for the establishment of a new mIGRA algorithm for interpretation. As such, the number of indeterminate cases was drastically reduced (80-100%) when the ConA + PWM mixture was used as a positive mitogen control along with a new mIGRA algorithm for interpretation.

## Introduction

With their close phylogenetic relationships and physiological similarities to humans, nonhuman primates (NHPs) have been commonly used to test the safety and efficacy of human pharmaceuticals [1-4]. Specified pathogen-free NHPs are highly demanded for these tests. Tuberculosis (TB)-free is an important requirement for imported laboratory NHPs, including cynomolgus macaque (*Macaca fascicularis*), for the Association of Primate Veterinarians (APV) and the Centers for Disease Control and Prevention in the United States and Canada [3, 5].

TB is a chronic bacterial infection caused by a group of *Mycobacterium tuberculosis* complex (MTBC), of which the species of greatest concern are *M. tuberculosis* (*M*.*tb*) and *M. bovis* [6, 7]. Aerosol transmission is the main factor in the outbreak of the disease. Like humans, after TB infection, cynomolgus macaque could be latent, subclinical, or active depending on the host’s immune system [8]. Therefore, early detection of TB is a primary requirement. A TB testing algorithm that incorporates multiple assays; for example, culture, molecular technique, antibody blood test, tuberculin skin test (TST), and interferon gamma release assay (IGRA), was suggested to improve overall sensitivity and specificity [9-11]. MTBC culture (directly detected MTBC) is supposed to be a gold standard method in humans, but it was inconvenient for routine screening in a large number of captive monkeys. Molecular techniques such as IS*6110* nested PCR or X-pert took a shorter time, but could not detect the immune response. Although the antibody blood test such as ELISA was developed, the humoral-mediated immune response appeared after the cell-mediated immune response, leading the TST and/or IGRA to be kept as preferable methods in many primate facilities [11] [12].

TST *in vivo* test is a classical tool for TB detection in NHP. The test uses purified protein derivatives (PPD) to stimulate host immune cells to secrete cytokines by intradermal injection to the eyelid. Swelling and redness 24, 48 and 72 hours after injection are monitored, meaning that at least two visits are needed. Other limitations of the TST are, for example, an expert demanding in steps of observation and interpretation, and a shortage of PPD. Thus, the IGRA is introduced. The IGRA blood test is convenient and does not require repeated handling of animals or extended intervals prior to retesting [13]. Two commercial IGRA kits, Quantiferon-plus (QFT-plus) and ELISpot kits, used two specific mycobacterial peptides of early secreted antigenic target 6 kDa (ESAT6) and 10 kDa culture filtrate antigen (CFP-10) to stimulate blood immune cells (mainly lymphocytes) and measured IFN-γ levels. Both ESAT6 and CFP10 are the most popular biomarkers due to their role in mycobacterial virulence and absence in the *M. bovis* BCG-vaccine strain [13-15]. Interpretation was carried out by comparing with negative (no mitogen added) and positive (mitogen added) controls. Recently, our team has introduced monkey IGRA (mIGRA) for the detection of TB in naturally TB infected macaques by combining human blood IGRA tubes (the QFT plus kit) and the monkey IFN-γ ELISA kit together [16]. However, many indeterminate results were unexpectedly encountered. Two factors were considered, high IFN-γ background level (NIL) and low mitogen (MIT) response. Aberrant IFN-γ expression of NIL without any stimulation is probably associated with autoinflammatory, autoimmune diseases, and viral infection of the hosts, which could not be directly correlated with TB infection ([17]. The low response to mitogen stimulation should be a mitogen of choice, such as a phytohemagglutinin (PHA) used in the QFT-plus kit.

Mitogens are nonspecific stimulants of immune cells that can activate various lymphocyte subpopulations to secrete cytokines such as IFN-γ [18, 19]. In the IGRA test, a mitogen serves as a positive control for the nonspecific T-cell response and reflects the response of active T lymphocytes in a host sample [20]. Commercially, commonly used mitogens are phytohemagglutinin (PHA), concanavalin A (Con A), and pokeweed (PWM). PHA, the lectin extract from the red kidney bean (*Phaseolus vulgaris*), contains potent, cell agglutinating and mitogenic activities [21]. PHA binds to the membranes of T cells and stimulates metabolic activity, cell agglutination, and mitogenic activities [22, 23]. Con A is another plant lectin (carbohydrate-binding protein) originally extracted from the jack bean (*Canavalia ensiformis*) and is known for its ability to stimulate mouse T-cell subsets, giving rise to four functionally distinct T cell populations, including precursors to regulatory T cells [24]. In addition, Con A does not have any mitogenic effect on B cells [25]. PWM, a lectin mitogen derived from the roots of *Phytolacca americana*, functions as a mitotic stimulus for the division of lymphocytes and specifically induces the proliferation of B cells, plasma cells, and T cells in mice and humans [26-28]. Since no report of the mitogenic effect and secreted IFN-γ levels of the IGRA test was found in cynomolgus macaques, this study was performed to compare 8 sets of mitogens on IFN-γ stimulation and to establish a new mIGRA algorithm for interpretation of TB infection.

## Materials and methods

### Animal ethics and permit

The experimental procedures in captive macaques at the Krabok-Koo Wildlife Rescue Center (KBK) were approved by the Institutional Animal Care and Use Committees of Mahidol University (Protocol review number: 009/2564) and the Department of Nationals, Wildlife and Plant Conservation of Thailand. This study was financially supported by NSRF through the Human Resources & Institutional Development, Research and Innovation Program Management Unit (grant number B05F640122) and partially supported by Drug Resistant Tuberculosis Fund under patronage of pass HRH Princess Galyanivadhana, Krom Loung Narathivas Rachnaharinth.

### Animals and blood collection

The subjects of this study were 316 cynomolgus macaques (*Macaca fascicularis*) reared in socially housed gang cages at KBK, Ta Takiap District, Chachoengsao Province, eastern Thailand, which is under the authority of the Department of the National Parks, Wildlife and Plant Conservation of Thailand. They were housed in the same cage or in the vicinity of the previously reported cynomolgus macaques naturally infected with *Mycobacterium tuberculosis* complex (MTBC) [16]. They were 250 (79%) males and 66 (21%) females ranging from 2 to 15 years of age with a body weight of 2-10 kg.

The monkeys were anesthetized with a mixture of zoletil (3–5 mg per kg) and dexmedetomidine hydrochloride (0.03–0.05 mg per kg), collected blood samples, and examined physical conditions. Blood was collected by femoral venipuncture, transferred to lithium-heparin tubes (cat. no. care-LIT4, Carestainer, China), and IFN-γ levels were determined using the two-step IGRA test (see details below). Blood samples were first collected from 12 monkeys for screening of 5 mitogen stimulations on IFN-γ levels. Twelve macaques (7 males and 5 females) aged 5.5-13 years with body weight of 3-11 kg were selected from 316 monkeys according to the criteria that they were negative for all TB tests, including TST, IGRA, ELISA (for ESAT6 and CFP10 antibody), and GeneXpert. One of the 5 mitogens that showed the highest IFN-γ level simulation was selected and tested again in all 316 monkeys.

Among 316 monkeys, 100 monkeys suspected of TB (70 individuals were tested positive for TST, IGRA, or GeneXpert, 14 individuals were in the same cage with a dead animal infected with MTBC, and the remaining 16 individuals were tested negative for TST, IGRA and GeneXpert) were selected and followed for 8 rounds at 0, 4, 7, 11, 14, 17, 20 and 26 months (M0, M4, M7, M11, M14, M17, M20 and M26, respectively) and used in the prospective study to determine mIGRA cutoff values.

### Mitogens

The mitogens used were phytohemagglutinin (PHA) from *Phaseolus vulgaris* (cat no. L1668, Sigma), Concanavalin A (Con A) from *Canavalia ensiformis* (cat no. C5275, Sigma), Lectin or pokeweed mitogen (PWM) from *Phytolacca americana* (cat no. L8777, Sigma). Commercially available phytohemagglutinin from the QuantiFERON-TB Gold-Plus kit (QFT-Plus; Catalog no. 622536, QIAGEN, USA) with unknown concentration in the mitogen tube (QFT-PHA) was also used as a reference.

### Comparison of mitogen stimulations on IFN-γ secretion

Among the 12 monkeys with negative TB selected, stimulations of 4 mitogen sets [PHA (20 μg/ml), Con A (20 μg/ml), PWM (20 μg/ml), Con A+PWM (20 μg/ml each in a ratio of 1: 1)] were compared to that of the QFT-PHA mitogen of the QFT-plus kit. Each 0.5 ml of blood was transferred to 4 mitogen tubes (PHA, Con A, PWM, and Con A+PWM) and 1.0 ml of blood was transferred to the QFT-PHA tube following the instruction of the QFT-plus kit within 16 hours after blood sampling. After stimulation, IFN-γ levels were determined by IFN-γ ELISA (see the details below) and optical density (OD) values at a wavelength of 450 nm were read. The OD_450_ value of the plasma background (OD_NIL_) was used to subtract the OD_450_ value of the mitogen (OD_MIT_), and the OD_MIT-NIL_ of the 5 mitogen sets was compared. Since the OD_MIT-NIL_ values of ConA+PWM was the highest (see Results), only the mitogen stimulation tests between ConA+PWM and QFT-PHA were repeated in all 316 monkeys.

### Plasma IFN-γ determination using the mIGRA test

In this study, the two-step mIGRA test; blood stimulation and measurement of IFN-γ level, was used. Five ml of blood was collected from each animal.

#### Blood stimulation

One ml of heparinized whole blood was aliquoted in each of the 5-format tubes (QFT-NIL, QFT-TB1, QFT-TB2, QFT-PHA and Con A+PWM) within 16 hours after blood sampling. All tubes were vertically tilted to mix antigens in the blood prior to incubation at 37°C for 20 hours. After incubation, plasma samples were harvested and stored at −80 °C before monkey IFN-γ ELISA assay (see below). All steps were carried out under the class II biosafety cabinet (BSC) (Model NU-440-600E, Nuaire, USA).

#### Determination of plasma IFN-γ Levels

Plasma sample was diluted with ELISA diluent (catalog number 3652-D2, Mabtech, Sweden) in a ratio of 1:2. The diluted plasma was determined to determine IFN-γ levels using a commercial monkey IFN-γ ELISApro kit (catalog no. M4210M-1HP-10, Mabtech AB, Sweden), containing mAb MT126L, mAb7-B6-1-biotin and human IFN-γ standards. Monkey IFN-γ levels were determined by measuring the OD at a wavelength of 450 nm (OD_450_). OD_450_ values of TB1, TB2, QFT-PHA, and Con A+PWM were subtracted with the OD_450_ value of the IFN-γ background (OD_NIL_) as OD_TB1-NIL_, OD_TB2-NIL,_ and OD_MIT-NIL_ before read-out. Before the interpretation of the mIGRA result, the cutoff values and the criteria were established as described in the result section.

### Statistical Analysis

The normal distribution of plasma IFN-γ levels was tested. If the normal distribution is detected, the IFN-γ levels are analyzed using one-way ANOVA with Tukey’s multiple comparison test. All statistical analyzes and graphical presentation were performed using GraphPad Prism 9 software version 9.0.0 (GraphPad, SanDiego, CA, USA). *p*-values less than 0.05 were considered significant differences.

## Results

### Comparison of mitogen stimulation on IFN-γ secretion of monkey blood cells

When comparing the OD_MIT-NIL_ of all 5 mitogen sets (PHA, Con A, PWM, Con A+PWM, and QFT-PHA) of 12 monkeys, it was found that the OD_MIT-NIL_ value of Con A+PWM was significantly higher than the other 4 mitogen stimulations (one-way ANOVA, *p* < 0.0001) as follows; Con A+PWM > PWM > QFT-PHA > Con A > PHA (**Fig 1 and S1 Table**). Monkey No. KBK114 showed low responses in all 5 mitogen stimulations (OD_450_ values ranging (-0.027) - 0.258), while Monkey No. KBK292 showed high responses in all 5 mitogen stimulations (OD_450_ values ranging 0.414 – 4.270). However, the stimulations of Con A+PWM remained higher than other simulations in these two monkeys. Note that 12 macaques gave higher OD_450_ values for QFT-PHA stimulation than for PHA (**S1 Table**).

**Fig 1:**
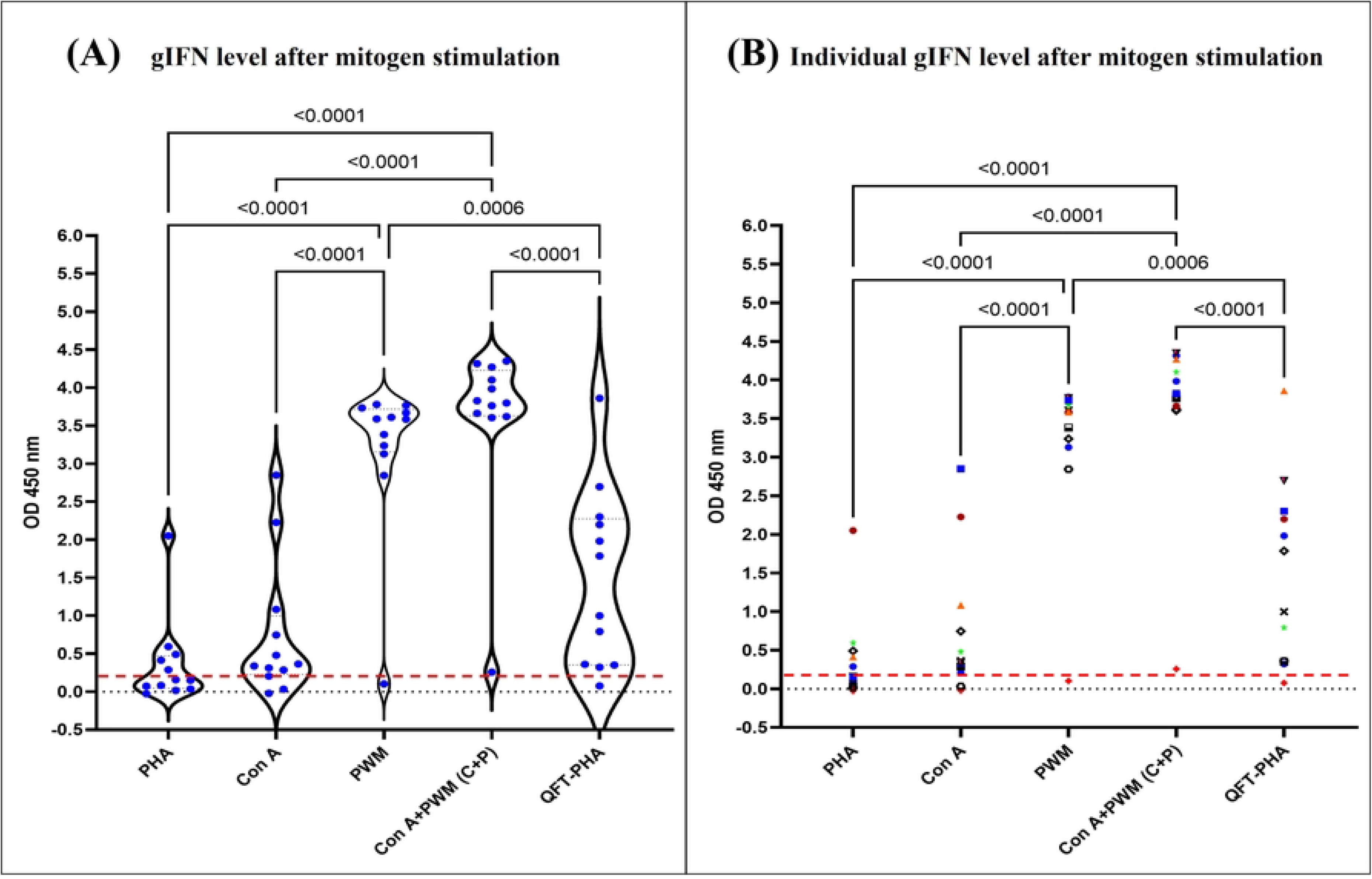
IFN-γ values as OD_MIT-NIL_, of 12 selected cynomolgus macaques after stimulation with prepared PHA (20 μg/ml), Con A (20 μg/ml), PWM (20 μg/ml), a mixture of Con A and PWM (C+P; 20 μg/ml each), and (QFT-PHA) and subtraction of the blank and plasma background. **Left panel:** The OD_450_ values of IFN-γ after mitogen stimulation for 20h at 37°C. All OD_450_ values are also marked as spots. **Right panel:** Individual OD_450_ values of IFN-γ levels responding to 5 mitogen sets and were obtained from 12 cynomolgus macaques. The red line indicated the 95th percentile of OD_NIL_ (= 0.18).

To confirm that mitogen selection for comparison with mIGRA results is one of the key factors to be considered, mitogen stimulation tests between Con A+PWM and QFT-PHA were repeated in all 316 cynomolgus macaques. Using QFT-PHA as a mitogen, monkeys showed OD_MIT-NIL_ values that ranged (-0.14) - 3.99 (mean±SD = 1.18±1.26). If the mixture of Con A+PWM was used as the mitogen, the OD_MIL-NIL_ values ranging by (-0.24) - 4.36 (mean±SD = 2.59±1.37) (**Fig 2**). Comparison of the IFN-γ response between QFT-PHA and Con A+PWM stimulation, 93.7% of the animals (296/316) showed the higher value of IFN-γ to the Con A+PWM mitogen than QFT-PHA (*p*-value < 0.0001), while 6% of the animals (19/316) showed the opposite direction. Surprisingly, 0.3% of the animals (1/316) did not show any response to any mitogen controls. Furthermore, it was found that 72 (22.8%) and 12 (3.8%) animals gave the OD_MIT-NIL_ values less or equal to 0.18 when stimulated with QFT-PHA or Con A+PWM, respectively. Moreover, there were 9 (2.8%) animals, giving the OD_NIL_ values higher than 0.18. Nevertheless, this suggests that Con A+PWM should be used as a mitogen to determine host immune status through plasma IFN-γ level after stimulation in cynomolgus monkeys. Thus, the mixture of Con A+PWM mitogen was selected for the next assays.

**Fig 2:**
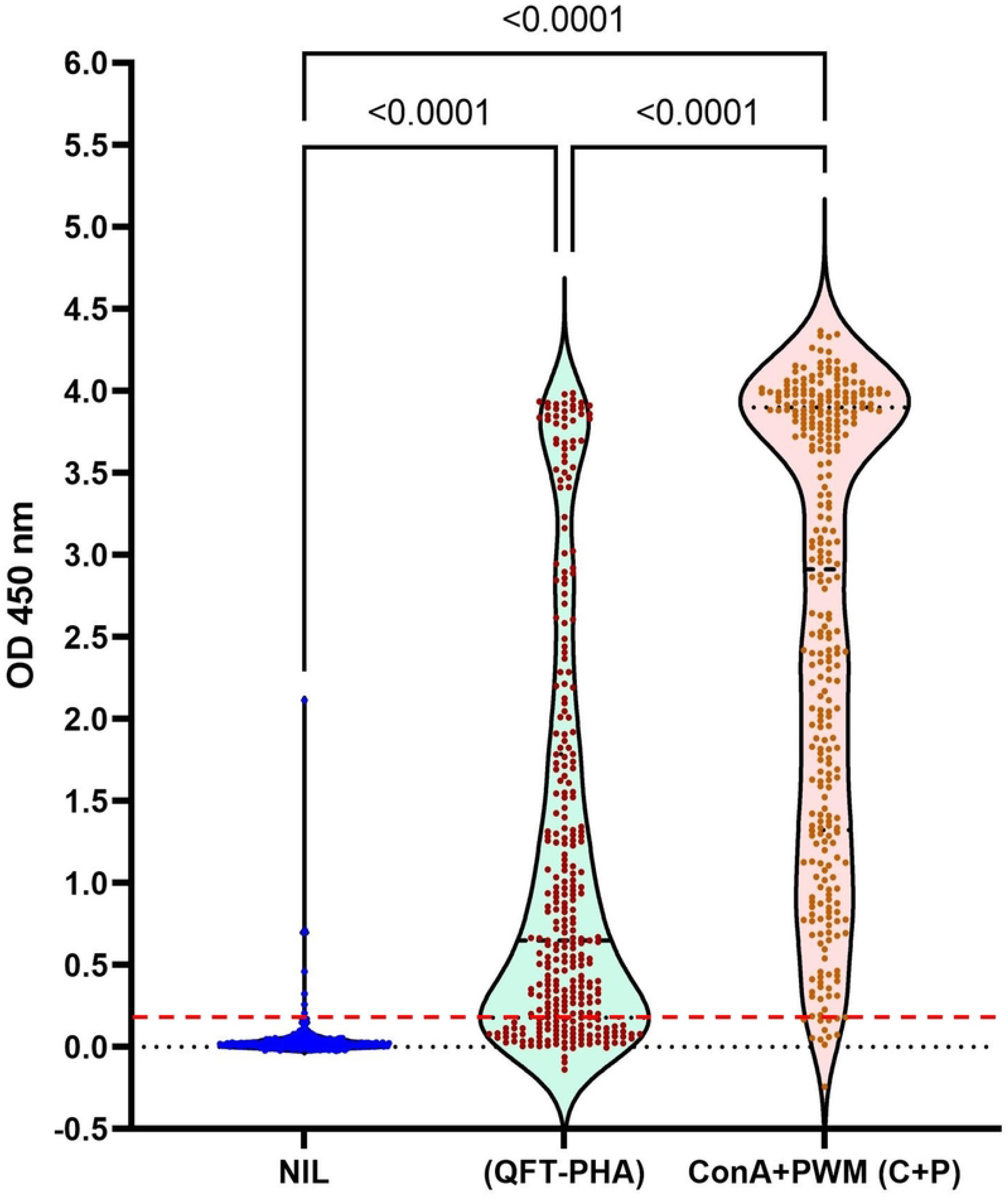
All IFN-γ levels as OD_450_ values obtained from whole blood immune cells from 316 naturally TB-contact cynomolgus macaques after stimulation with QFT-PHA and Con A+ PWM (C+P) mitogens and without any stimulator (The NIL tube). All minimum and maximum data values were plotted together with the mean (+) and median (a line) values in a box. The red line indicated the 95th percentile of OD_NIL_ (= 0.18).

### Determination of mIGRA cutoff values and a new algorithm to report TB Λ

The IFN-γ OD_450_ values were obtained from 5-format tubes (NIL, TB1, TB2, QFT-PHA and Con A+PWM), calculated and presented as OD_NIL_, OD_TB1-NIL_, OD_TB2-NIL,_ OD_(QFT-PHA) -NIL_ and OD_(ConA+PWM)-NIL_ values. Using of standards in a concentration of 7.87, 15.75, 31,25 and 62.5 pg/ml according to the monkey γ-IFN ELISA kit manual (Cat. No. 3421M-1H-20, Mabtech, Sweden), the percentage of the inter-assay and intra-assay coefficient of variation (%CV) were determined and found to be in a range of 23-26% and 0-15%, respectively. To determine the cutoff value, the OD_450_ values obtained from 100 selected cynomolgus macaques, which were followed for 8 rounds (M0, M4, M7, M11, M14, M17, M20 and M26), total 721 values, were pooled and analyzed. Using a box plot analysis, the 95^th^ percentile of OD_NIL_ was calculated and found to be 0.18. Next, the OD_450_ values of TB1 and TB2 that were higher than the OD_NIL_ value (or OD_TB1-NIL_ and OD_TB2-NIL_ > OD_NIL_) were selected. The 25% (Q1) and 75% (Q3) percentile values of OD_TB1-NIL_ and OD_TB2-NIL_ and the interquartile range (IQR) values were calculated. Outlier values were identified from an upper limit value [Q3+ (1.5*IQR)] and a lower limit value [Q1-(1.5*IQR)]. As a result, the upper and lower limit values of both OD_TB1-NIL_ and OD_TB2-NIL_ were 0.05 and -0.04, respectively.

Following the QFT-plus kit handout, it suggested that after stimulation, the level of IFN-γ of the TB tubes (TB1 or TB2) should be higher than 25% of the individual IFN-γ value of the NIL tube (OD_NIL_); thus we included this value as an additional cutoff value in our study. Therefore, OD_TB1-NIL_ and OD_TB2-NIL_ were determined from the two cutoff values, 0.05 and 25% of the individual OD_NIL_ value (**Fig 3**). With respect to these criteria, the interpretation of the IFN-γ ELISA is as follows. If the OD_NIL_ value was ≤0.18, OD_MIT-NIL_ > OD_NIL,_ and the OD_TB-NIL_ was ≥0.05 and ≥25% of individual OD_NIL_, the mIGRA result was interpreted as “positive”. If the OD_NIL_ value was ≤0.18, OD_MIT-NIL_ > OD_NIL_, and the OD_TB-NIL_ was <0.05, the mIGRA result was interpreted as “negative”. If the OD_NIL_ value was >0.18, indicating the high IFN-γ background. Thus, IFN-γ secretion after stimulation with TB1 or TB2 was invalid, and the mIGRA result was interpreted as ‘indeterminate’. Furthermore, if the OD of mitogen-positive controls [OD_(QFT-PHA)_ and OD_(ConA+PWM)_] were lower or equal to the OD_NIL_, the mIGRA result was also interpreted as ‘indeterminate’. This indicated that host immune cells were inactive or impaired host cell-mediated immunity.

**Fig 3:**
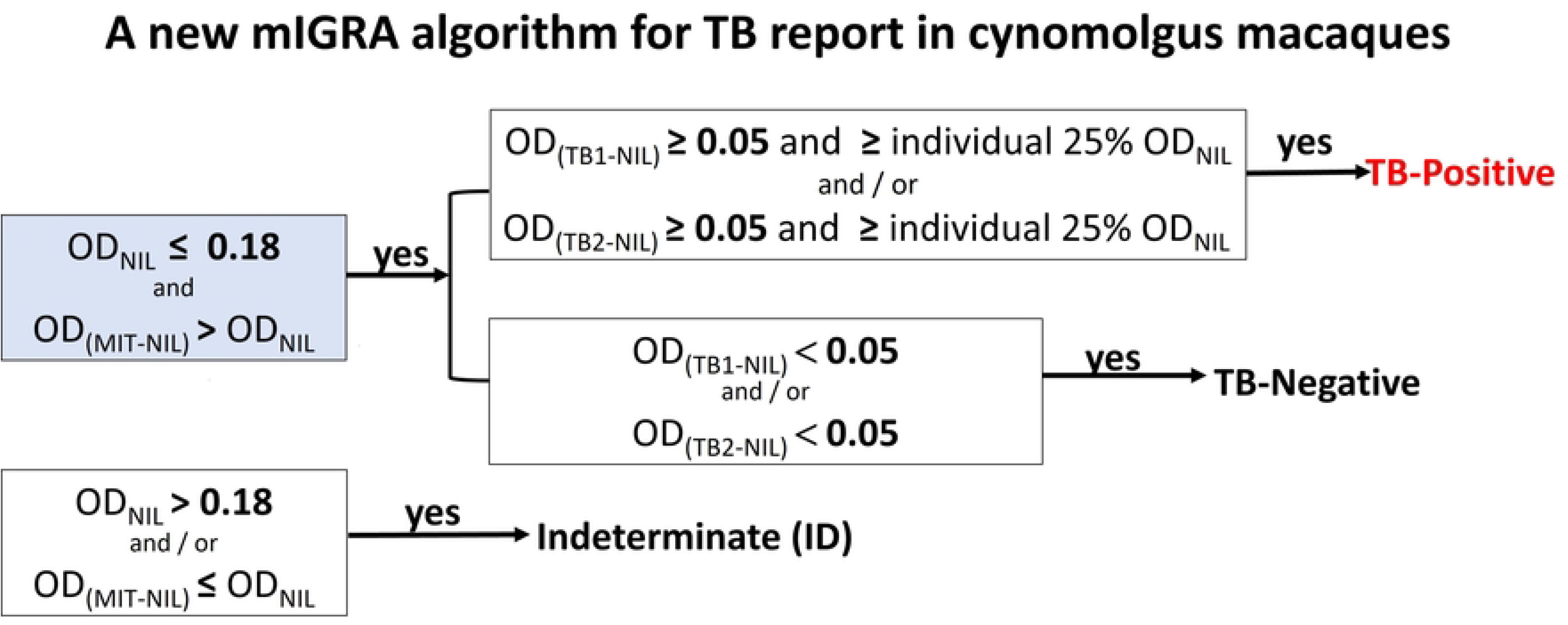
A flow chart of the new mIGRA algorithm for TB report in cynomolgus macaques using QFT plus tubes (NIL, TB1 and TB2) and a mixture of Con A and Pokeweed mitogen (Con A+PWM) as a positive mitogen control. OD is an optical density reading at wavelength of 450 nm. The TB1 tube contains ESAT6 and CFP10 peptides specific to CD4+-T cells, whereas the TB2 tube contains ESAT6 and CFP10 peptides specific to CD4+ and CD8+ -T cells. The NIL tube means that there is no mitogen stimulation. The MIT tube contains either a mitogen ConA+PWM or QFT-PHA, which was applied in this study.

Together, a new algorithm for the interpretation of mIGRA was proposed using the OD_450_ values (**Fig 3**). After considering OD_NIL_ ≤ 0.18 and OD_MIT-NIL_ > OD_NIL_, if the OD_TB1/2-NIL_ was ≥0.05 and ≥25% of the individual OD_NIL_ value, the positive was reported and likely implied the MTBC infection. If OD_TB1/2-NIL_ was <0.05, the negative was reported and likely implying non-MTBC infection. However, indeterminate (ID) was reported, if either OD_MIT-NIL_ ≤OD_NIL_ or OD_NIL_>0.18 was detected.

### Comparison of TB infection result between the use of QFT-PHA and Con A+PWM mitogens

After a new mIGRA algorithm was identified as mentioned above, the interpretation of the TB infection results of 100 cynomolgus monkeys was performed. The use of QFT-PHA and Con A+PWM as positive mitogens was compared. As shown in **Table 1**, the positive cases of mIGRA were the same regardless of comparison with any mitogen type, that is 17, 16, 13, 12, 14, 15, 13, and 8 positive cases for M0, M4, M7, M11, M14, M17, M20 and M26, respectively. To prevent the outbreak, after M4, the positive cases were separated from their gang cages and moved to a clean remote area for further investigation and management. However, 3 positive mIGRA cases were later found death during M9-M10. Their lesions in the lungs and liver organs were inspected and the infection with *Mycobacterium tuberculosis* was confirmed by the culture method.

**Table 1:** TB interpretation with the new algorithm of mIGRA using either QFT-PHA or Con A + PWM as positive mitogens. Data were obtained from the 8-round prospective study of 100 naturally TB-contacted cynomolgus macaques in month-0 (M0), month-4 (M4), month-7 (M7), month-11 (M11), month-14 (M14), month-17 (M17), month-10 (M20), and month-26 (M26). At 7 rounds in M4, M7, M11, M14, M17, M20 and M26, blood collection was not applicable, or animal loss, or animal death occurred; thus, the total cases were less than 100 animals. The asterisk (*) indicates 3 positive cases infected with *M*.*tb*, which died in M9-M10.

| mIGRA interpretation |  | QFT-PHA mitogen (Cases) |  |  |  |  |  |  |  | Con A+PWM mitogen (Cases) |  |  |  |  |  |  |  |
| --- | --- | --- | --- | --- | --- | --- | --- | --- | --- | --- | --- | --- | --- | --- | --- | --- | --- |
|  |  | M0 | M4 | M7 | M11 | M14 | M17 | M20 | M26 | M0 | M4 | M7 | M11 | M14 | M17 | M20 | M26 |
| Positive |  | 17 | 16* | 13* | 12 | 14 | 15 | 13 | 8 | 17 | 16* | 13* | 12 | 14 | 15 | 13 | 8 |
| Negative |  | 50 | 66 | 68 | 29 | 51 | 49 | 59 | 46 | 76 | 72 | 79 | 63 | 66 | 65 | 65 | 69 |
| Indeterminate | OD<sub>NIL</sub> value more than 0.18 (95percentile of OD<sub>NIL</sub> value) | 1 | 9 | 3 | 4 | 7 | 6 | 4 | 7 | 1 | 9 | 3 | 4 | 7 | 6 | 4 | 7 |
|  | OD<sub>(MIT-NIL)</sub> value less or equal to 0.18 | 32 | 7 | 12 | 43 | 15 | 16 | 6 | 23 | 6 | 1 | 1 | 9 | 0 | 0 | 0 | 0 |
| Total (Cases) |  | 100 | 98 | 96 | 88 | 87 | 86 | 82 | 84 | 100 | 98 | 96 | 88 | 87 | 86 | 82 | 84 |

When the use of QFT-PHA and Con A + PWM mitogens was compared, the mIGRA results were markedly different in indeterminate cases. The lower indeterminate cases were detected when the Con A+PWM mitogens were used compared to that of the QFT-PHA (**Table 1**). The ratio of the indeterminate cases of QFT-PHA: ConA+PWM were 33:7, 16:10, 15:4, 47:13, 22:7, 22:6, 10:4, and 30:7 for M0, M4, M7, M11, M14, M17, M20 and M26, respectively. Moreover, if classified the ID cases into 2 categories as mentioned in the new mIGRA algorithm, it could be presented that (1) ID causing from OD_NIL_>0.18 of QFT-PHA: ConA+PWM were in the same ratio as 1:1, 9:9, 3:3, 4:4, 7:7, 6:6, 4;4 and 7:7 for M0, M4, M7, M11, M14, M17, M20 and M26, respectively. While (2) ID causing from OD_MIT-NIL_ ≤OD_NIL_0.18 of QFT-PHA: ConA+PWM were 32:6, 7:1, 12:1, 43:9, 15:0, 16:0, 6:0 and 23:0 for M0, M4, M7, M11, M14, M17, M20 and M26, respectively. Aligned with the indeterminate cases, the ratio of the negative cases of QFT-PHA: Con A+PWM were 50:76, 66:72, 68:79, 29:63, 51:66, 49:65, 59:65 and 46:69 for M0, M4, M7, M11, M14, M17, M20 and M26, respectively. Combining the use of ConA+PWM mitogen and a new algorithm of mIGRA interpretation, 80-100% of the indeterminate results were reduced.

## Discussion

With the high demand for nonclinical drug and vaccine tests, especially during the COVID-19 pandemic, the specific pathogen-free (SPF) cynomolgus macaques were required. TB is designated as the first infectious disease that must be screened and infected macaques must be removed from the colony immediately. Among various TB detection tools, IGRA is acceptable as a method to detect early TB infection and the latent period [11, 12]. As such, the monkey IGRA assay was developed [16]. A major hindrance of the mIGRA assay in monkeys is a highly indeterminate result because of the non-stimulation of the mitogen used in the assay. The IGRA mitogen is a very important reference for the comparison and interpretation of the results. If the mitogen-stimulated value passes the cutoff value, it can ensure the host immune status for the IGRA interpretation. In humans, the low mitogen response found in the IGRA test is most often associated with reduced lymphocyte activity due to insufficient lymphocytes, incorrect addition of the mitogen, prolonged specimen transport, improper specimen handling, or the presence of antibodies to IFN-γ [29-32]. In NHPs, two commercial TB-IGRA kits, PrimaGAM and gamma-interferon test (GIFT) assay, used different mitogens to stimulate lymphocytes. In PrimaGAM, Con A was used as a positive control mitogen [33], while Con A+PWM was used as a positive control in the GIFT assay [34]. However, the concentration and minimal response to Con A were not determined. Apart from Con A and PWM, PHA was also used as a mitogen for IGRA and inappropriate stimulation of immune cells of cynomolgus monkeys and chacma baboons was reported [30, 35]. Until now, no suitable mitogen for the IGRA test in cynomolgus monkeys has been widely recommended. Since MTBC infection can result in immunosuppression, stimulation with the appropriate mitogens should be used to ensure that the animal is immune competent enough to rule out a false negative response for the IGRA test. Here, we compared 4 sets of prepared mitogens and one commercially available QFT-PHA mitogen in the mIGRA assay of cynomolgus macaques. We also proposed a new algorithm of TB interpretation, based on mIGRA levels determined using mIGRA and selected mitogen, in cynomolgus monkeys.

Although mitogen stimulation results depended highly on the concentration used and the varied doses (dose-response) of mitogen were suggested in the test [20, 36], the high blood volume needed from each animal is a limitation. In this study, we selected 4 mitogens in a fixed dose of 20 μg/ml for the stimulation of IFN-γ release from T cells of cynomolgus monkeys, and the results were ranked by Con A+PWM > PWM > QFT-PHA > Con A > PHA. Our results aligned with the previous study that the QFT-PHA mitogen could activate higher IFN-γ secretion from the whole blood of cynomolgus macaques than Con A [20]. It should be noted that the concentration of Con A used in the previous study (5 μg/ml) was lower than our study (20 μg/ml). Compared to the previous study in human PBMC isolated from healthy donors, the same order of stimulation was found, although the concentrations of mitogens were lower than in our study; PWM (5 μg/ml) > PHA (10 μg/ml) > Con A (5μg/ml) [25]. A slight difference was observed in other human whole blood derived from healthy volunteers, and a preferential order was PWM (2 μg/ml) ≥ Con A (5 μg/ml) ≥ QFT-GIT-PHA (the PHA mitogen from the Quantiferon Gold kit) > PHA-P (10 μg/ml) [20, 37]. This implies that mitogen stimulation results depend on the concentration used, cell density, and culture period [36]. If a constant number of lymphocytes is used, the dose-response sigmoid curve is usually presented. The stimulation decreases at the very high mitogen concentration, which exceeds the optimal concentration, presumably due to nonspecific toxicity to the cells. Focusing on the PHA mitogens used in our study, the QFT-PHA mitogen stimulated IFN-γ secretion from the whole blood of cynomolgus macaques stronger than the prepared PHA (20 μg/ml). This suggested that the concentration of PHA used in the QFT kit could be greater than 20 μg/ml. Similarly to the report in human whole blood, the commercial QFT-GIT-PHA mitogen gave a higher IFN-γ level than that of the prepared PHA-P mitogen (20 μg/ml) [20]. The prepared PHA in our study was also the PHA-P form, while those in the QFT-GIT or QFT-plus kit were unknown. Commercially, two forms of PHA are sold, the PHA-P protein form and the PHA-M mucoprotein form. We suspected that the higher IFN-γ response after QFT-PHA stimulation compared to prepared PHA stimulation could be either a higher dose of mitogen or a different form of PHA used.

The different actions of mitogens to stimulate human immune cells were caused by the difference in specific noncatalytic carbohydrate recognition domains (or glycoproteins) in their biochemical/physicochemical properties, carbohydrate binding specificity, and biological activities. The specific glycol-conjugated domain of mitogens binds to its primary target in the carbohydrate moiety of cell membrane glycoproteins known as the Toll-like receptor (TLR). Con A binds to α-D-mannosyl residue, while PWM and PHA bind to N-acetyl-D-glucosamine [38]. In humans, PHA stimulates TLRs (-2/6, -4, and -5) on CD^3+^/CD^8+^ T cells. Con A binds to TLRs (-2/6) on CD^4+^/CD^8+^ T cell and shows higher mitogenic activity in CD^4+^ than the CD^8+^, resulting in enhanced proliferation and IFN-γ production [25]. PWM not only stimulate the TLRs (-2/6, -4, and -5) in CD3 + / CD4 + / CD8 + T cells, but also contains signals from TLR4 proteins for the B cell, leading to its effect on both the T cell and the B cell for a proliferative response and produced IFN-γ [39]. This should be one of the reasons why PWM had stronger mitogenic activity and stability compared to PHA and Con A in our study. The combination of Con A and PWM might enhance the number of specific TLR on the surface of immune cells to be stimulated, and thus the level of IFN-γ secretion after the Con A+PWM stimulation was higher than a single mitogen used. No indeterminate result was encountered when a mixture of Con A and PWM was used as the mitogen positive control compared to a single used mitogen (2 indeterminate cases for Con A and 1 indeterminate case for PWM).

In addition to mitogen type, heat or cation-supplemented mitogen (Mg^2+^, Ca^2+^ and Zn^2+^) could affect IFN-γ production [20]. If the animals are supplemented with cation such as calcium, detection of tuberculosis using the IGRA test should be considered. As seen in **Fig 1**, IFN-γ production in response to each mitogen in each monkey was individually distinctive; some showed high response, and some showed low response, which depends on host immunity status. Therefore, it should be taken as awareness during diagnosis.

It needs to be kept in mind that the different species of animals tested, the response to mitogen stimulation might be different. For example, PWM and PHA showed the same activity in inducing IFN-γ secretion from the whole blood cells of wild dogs [40]. Different positive control mitogens were suggested, that is, PWM for domestic cattle (*Bos taurus*) [41]; Con A for Asian elephants (*Elephas maximus*) [42]; and PHA for human (QFT-plus, QIAGEN). In addition to species of animals, different genetic inheritances between different individuals in the same cohort should also be considered. When comparing IFN-γ secretion between QFT-PHA and Con A+PWM stimulation, 93% of cynomolgus macaques showed a higher level of IFN-γ after ConA + PWM stimulation than QFT-PHA, while it was opposite for 5% of animals. We hypothesized that the host’s genetic profile might be different, leading to different specific membrane glycoproteins on immune cells. Therefore, the species of animals, genetic background, sample type (whole blood or PBMC), and the IGRA protocol (such as the duration time of blood collection before the test, blood handling, and a period of stimulation) should be taken into account when you perform the IGRA test.

When changing the mitogen positive control used from QFT-PHA to Con A+PWM together with a new mIGRA algorithm to determine TB status in a large cohort (100 monkeys, 8 rounds), the number of indeterminate cases in a condition of OD_MIT-NIL_ ≤OD_NIL_ 0.18 was significantly reduced (80-100%). Unfortunately, although the mitogen was changed, the number of indeterminate, causing from OD_NIL_>0.18, was remaining and be the same. This implied that the latter ID group of animals would have healthy problem or inflammation leading to have high γ-IFN background and should be taken into accounts for follow-up. Therefore, this suggests that the Con A+PWM mitogen should be applied to mIGRA for the detection of tuberculosis in cynomolgus monkeys. Due to the smaller number of indeterminate cases, the small number of animals would pass through the euthanasia process. However, the indeterminate cases remained in the Con A+PWM group. The explanation could be (1) an insufficient immune response to the mitogen due to the impaired host immune system, that is, suspected immunosuppression, chronic disease, young age, and malnutrition, (2) technical errors (delay in sample delivery or vigorous handling) during sample collection, or (3) anergy. When the indeterminate cases are detected by mIGRA, it is likely that the host is MTBC-infected, those cynomolgus macaques should be followed, and other assays should be ancillary used. Several TB detection methods, based on the cellular immune response as seen in our mIGRA, were GIFT, T-Spot, or ELISpot, and TST. They differ in the form of the test (*in vitro* whole blood test for mIGRA and GIFT; *in vitro* PMBC test for T-Spot and ELISpot; *in vivo* test for TST), the antigens used (specific TB peptide antigens for mIGRA; bovine-PPD and avian-PPD for GIFT, and MOT-PPD for TST), and the criteria for interpretation [9], [13], [16], [34]. Thus, the test results might not be interchangeable [31].

## Acknowledgments

The authors would like to thank all staffs at Krabokkoo Wildlife Breeding Center for technical assistance, with the permission of the Department of the National Parks, Wildlife and Plant Conservation, Thailand.

## Authors’ Contributions

SW, SM, PS, TPr, TPa, SM and AC: participated and study design. SW, SW, PC, NM, PS, MK, KN, AP, TK, RK, PP, and SM: - participated in the collection, testing, and interpretation of the data. SW, TPa, SM and AC: Writing – original draft, conceptualization, methodology, and formal analysis. SW, SM, PS and AC: review & editing, and project administration. All authors reviewed and approved the final manuscript.

## Supporting information

**S1 Table:** The IFN-γ values of individual 12 selected cynomolgus macaques after the subtraction of plasma and IFN-γ (NIL) background, as OD_MIT-NIL_. All values were obtained after stimulation with prepared PHA (20 μg/ml), Con A (20 μg/ml), PWM (20 μg/ml), Con A+PWM (20 μg/ml each), and (QFT-PHA). Red box indicated the values below the 95^th^ percentile of the OD_NIL_ (= 0.18).

